# Ultraviolet and yellow reflectance but not fluorescence is important for visual discrimination of conspecifics by *Heliconius erato*

**DOI:** 10.1101/088781

**Authors:** Susan D. Finkbeiner, Dmitry A. Fishman, Daniel Osorio, Adriana D. Briscoe

## Abstract

Toxic *Heliconius* butterflies have yellow hindwing bars that – unlike their closest relatives – reflect ultraviolet (UV) and long wavelength light, and also fluoresce. The pigment in the yellow scales is 3-hydroxy-DL-kynurenine (3-OHK), found also in the hair and scales of a variety of animals. In other butterflies including pierids, which similarly display wing colors that vary in both the UV and the human-visible range, behavioral experiments have indicated that only the UV component is most relevant to mate choice. Whether in *Heliconius* butterflies it is the UV, the human-visible yellow, and/or the fluorescent component of yellow wing coloration that is relevant to mate choice is unknown. In field studies with butterfly paper models we show that both UV and 3-OHK yellow act as signals for *H. erato* but attack rates by birds do not differ significantly between the models. Furthermore, measurement of the quantum yield and reflectance spectra of 3-OHK indicates that fluorescence does not contribute to the visual signal under broad-spectrum illumination. Our results suggest that the use of 3-OHK pigmentation instead of ancestral yellow was driven by sexual selection rather than predation.

**Summary statement:** *Heliconius* butterflies use a co-opted yellow pigment for communication, while predators are fooled by non-*Heliconius* mimics using ancestral yellow pigments.

## Introduction

Color patches of animals are complex traits composed of multiple components (Grether et al., 2004). The pigment cells known as chromatophores in the skin of fishes, reptiles and amphibians for example are color-generating structures comprised of distinct pigmentary and structural layers that vary in their ability to reflect light. The feather barbs or integument of birds or the wing scales of butterflies similarly have diverse nano-structure architectures, thin films, and pigments, which produce a dazzling array of colors (Prum and Torres, 2003; Vukusic and Sambles, 2003; Shawkey and Hill, 2005; Stavenga et al., 2011, 2014). These pigmentary and structural components of color patches work in tandem to produce signals used in a variety of contexts (e.g., crypsis, mimicry, aposematism, and mate choice). Since the biochemical and developmental mechanisms underlying pigmentary and structural properties of color differ, each of these components may be subject to different selective pressures, and hence independent evolutionary trajectories (Grether et al., 2004). Here we look specifically at how two components of a butterfly visual display, UV reflectance and human-visible yellow reflectance due to selective filtering by a specific wing pigment, may function as a signal in mate choice and predation. We also look at what contribution fluorescence makes, if any, to the signal.

Many butterfly species have colorful wing patterns in both the human-visible (400-700 nm) and in the UV (300-400 nm) ranges (Silberglied and Taylor, 1978; Eguchi and Meyer-Rochow, 1983; Meyer-Rochow, 1991; Rutowski et al., 2005; Briscoe et al., 2010). While the idea that UV coloration—invisible to humans—may serve as a ‘private channel’ of communication has been challenged (Cronin and Bok, 2016; but see Cummings et al., 2003), there is ample evidence that UV signals are important in animal communication (Rutowski, 1977; Johnsen et al., 1998; Smith et al., 2002; Cummings et al., 2003; Robertson and Monteiro, 2005; Kemp, 2008; Obara et al., 2008; Detto and Blackwell, 2009; Painting et al., 2016). On the other hand, although many butterflies have UV-visible color patches, in the absence of behavioral evidence, it is unclear whether the UV reflectance functions as a signal, or if it is simply an epi-phenomenon of the scale structure overlaying pigment granules. The same question can of course be applied to the colors produced by the pigments.

Studies of several butterfly groups suggest in fact that for color patches with both UV and visible reflectance, only variation in the UV component of the signal affects mate choice. Pierid butterfly males, *Colias eurytheme* and *C. philodice,* have forewing colors with both UV-iridescence due to the structural scattering of light by the scale lamellae (Ghiradella, 1974) and yellow-orange due to pterin pigments (Watt, 1964). In behavioral experiments, female *Colias* were shown to use the UV-reflection difference between the two species as a mate and species recognition cue, but not the human-visible color difference (Silberglied and Taylor, 1978). Female *Eurema hecuba* (Coliadinae: Pieridae) were similarly shown to prefer males with the brightest UV iridescence overlaying a diffuse pigment-based yellow (Kemp, 2007a). Given that many other butterflies have color patches with UV-visible reflectances, and that butterfly color vision systems are astonishingly diverse (Arikawa et al., 2005; Briscoe and Bernard, 2005; Stalleicken et al., 2006; Koshitaka et al., 2008; Sison-Mangus et al., 2008; Chen et al., 2013), it is worthwhile to investigate in other species whether it is the UV or the human-visible or both parts of the color patch reflectance spectrum that is being used for signaling. It is particularly interesting to investigate this question where there has been a phylogenetic transition from using one type of pigmentation to another, as for the yellow wing colors in the passion-vine butterflies of the genus *Heliconius* (Briscoe et al., 2010; Bybee et al., 2012)(see below).

*Heliconius erato* has yellow scales on its hindwings that contain the pigment 3-hydroxy-DL-kynurenine (3-OHK) (Tokuyama et al., 1967; Reed et al., 2008). The yellow bars reflect UV light and have a step-like reflectance at longer wavelengths ‐a rapid rise then a plateau in reflectance in the visible (400-700 nm) range (yellow lines, Fig. 1A,B)(see also Stavenga et al., 2004). Either the UV or the human-visible part of 3-OHK wing reflectance or both may serve as a signal for inter‐ and intra-specific communication. Intriguingly, 3-OHK’s appearance in *Heliconius* co-occurred with the evolution of the butterflies’ duplicated UV opsins, UV1 and UV2 (Briscoe et al., 2010; Yuan et al., 2010; Bybee et al., 2012). In some *Heliconius* species, UV1 and UV2 are found in both males and females (McCulloch and Briscoe, unpublished data). In *H. erato,* UV1 is a female-specific UV receptor with *λ*_max_=355 nm, while UV2 is a violet receptor with *λ*_max_=390 nm found in both sexes (Fig. 1, triangles and x marks, respectively) (McCulloch et al., 2016).

**Fig. 1.**
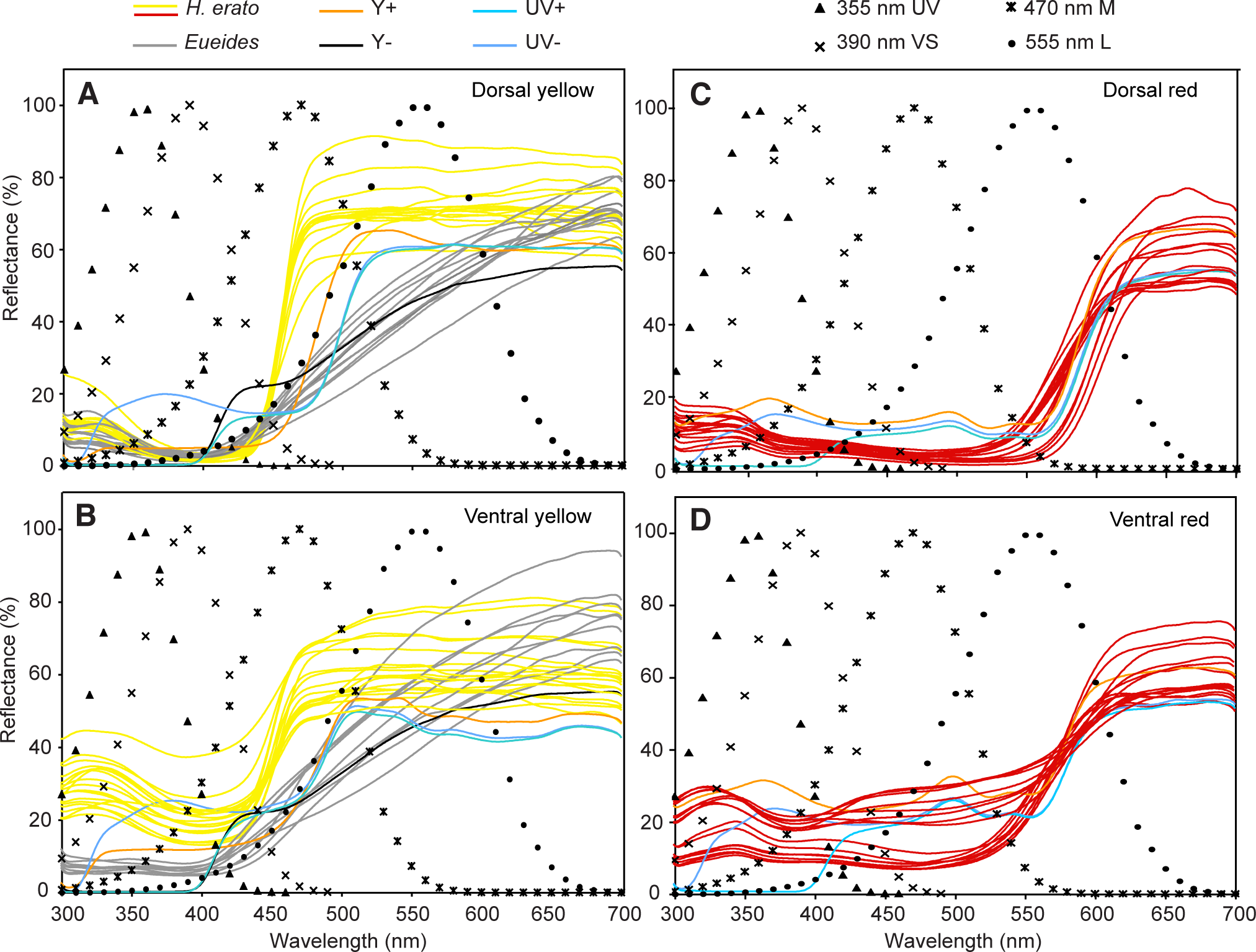
Reflectance spectra of *Heliconius erato* and *Eueides* wing colors and paper model colors used in the mate choice and predation experiments. (A) Dorsal yellow, (B) ventral yellow, (C) dorsal red, (D) ventral red. Reflectance spectra correspond to: *H. erato* (yellow or red), *Eueides* (grey), Y+ (orange), Y− (black), UV+ (dark blue), UV- (blue-green). Black symbols correspond to the spectral sensitivities of *H. erato* photoreceptor cells with peak sensitivities *−*_max_ at: 360 nm (triangles), 390 nm (×), 470 nm (*), and 555 nm (filled circles) (McCulloch et al., 2016). The photoreceptor with a peak at 360 nm is found in female but not male *H. erato*.

In addition to the components of the 3-OHK visual signal mentioned above, the yellow wing bars of *Heliconius* fluoresce under a hand-held blacklight (Movie S1). Fluorescence occurs when short-wavelength light is absorbed and then re-emitted as a longer wavelength, i.e. lower energy light. Fluorescent pigments are widespread in nature (Vukusic and Hooper, 2005; Lagorio et al., 2015) and are typically identified using spectrally narrow-band light; however, terrestrial illumination has a broad spectrum so it is unclear whether or not a pigment’s fluorescence contributes much to a potential signal under natural conditions. The emission spectra of the 3-OHK pigment overlaps with the visible part of the reflectance spectrum of 3-OHK on *Heliconius* wings (see below) and so would be well-suited to being detected by the blue-sensitive receptor of *H. erato* with *λ* =470 nm if it did (McCulloch et al., 2016).

Butterflies from the genus *Eueides,* which are a sister taxon to *Heliconius*, have mimetic wing patterns strikingly similar to some *Heliconius* species. These two genera co-occur in the same habitats, yet their yellow wing pigments lack the step-like reflectance spectrum of 3-OHK (grey line, Fig. 1A,B) (Bybee et al., 2012), and they do not fluoresce (data not shown). The yellow pigments in both butterflies appear similar to the human eye in natural light, but their spectra differ strongly (yellow and grey lines, Fig. 1A,B). Although modeling of wing colors suggests in principle that *Heliconius* can distinguish between *Heliconius* 3-OHK yellow and *Eueides* yellow (Bybee et al., 2012), it remains unknown whether *Heliconius* actually do so in nature. Previous work has shown that *H. erato* prefer chromatic over achromatic signals in the context of mate choice (Fig. S1) (Finkbeiner et al., 2014); but it is unclear whether the visible, the UV, or both parts of the reflectance spectrum of 3-OHK and fluorescence contribute to signaling. Prior work has also shown that avian predators will differentially attack achromatic compared to chromatic butterfly paper models (Fig. S1) (Finkbeiner et al., 2014; Dell’Aglio et al., 2016), but it is unknown whether avian predators will differentially attack butterfly paper models that vary in yellow coloration resembling the differences between *Heliconius* and *Eueides* yellow. While *Heliconius* wing color patterns warn avian predators of their toxicity (Benson, 1972; Chai, 1986), 3-OHK may further serve as a conspecific signal especially in courtship (Bybee et al., 2012; Llaurens et al., 2014). Demonstrating that *Heliconius* species can in fact discriminate 3-OHK yellow from other yellows in nature is an important step in elucidating the adaptive significance of 3-OHK pigmentation.

To further investigate the contribution of 3-OHK to *Heliconius erato* signaling, we carried out two sets of experiments: The first set tested responses of both male and female *H. erato* to four types of colored models, whose spectra were intended to approximate either those of *Heliconius* species or their mimics, such as *Eueides.* The first pair of spectra, which are designated Y+ or Y−, resemble 3-OHK (*Heliconius*) yellow or *Eueides* yellow. The second set of reflectance spectra have identical yellow and red coloration in the visible range, but UV reflectance is either present (UV+) or absent (UV-).

The second, complementary set of experiments tests the hypothesis that predatory birds will not differentially attack 3-OHK yellow from other yellows when presented with model butterflies due to the aposematic function of yellow in general. Together these experiments substantiate and elaborate our understanding of the function of 3-OHK yellow and UV coloration. We show also that fluorescence – although clearly visible in laboratory conditions, but with illumination restricted to the UV excitation wavelengths – is not likely to have any impact under the broadband and relatively low UV illumination found in nature.

## Material and methods

### Butterfly Models, Wing Reflectance Spectra, Environmental Light and Discriminability

Four paper model types of the *Heliconius erato petiverana* butterfly were made as described in Finkbeiner et al. (2012) with their colors modified as follows: with (Y+) and without (Y−) 3-OHK yellow, and with (UV+) and without (UV-) ultraviolet reflectance. The Y+ treatment had 3-OHK on the yellow portion of the wing (0.010 mg/¼l and 0.015 mg/¼l 3-OHK in methanol applied to the ventral and dorsal sides, respectively). This provided the models with the same pigment as found in the butterfly yellow scales. The yellow portion of the non-3-OHK yellow models (Y−) was covered with yellow Manila paper (Creatology^®^ Manila Drawing Paper, Item No. 410590). Manila paper has a reflectance spectra that resembles non-3-OHK yellow reflectance from the sister-genus to *Heliconius, Eueides,* which is a *Heliconius* mimic (Bybee et al., 2012) (Fig. 1A,B, grey and black lines). A thin film UV filter (Edmund Optics, Item No. 39-426) was placed over the Manila paper to create a closer match to *Eueides* yellow pigment. As a control, Mylar film was added to the yellow portions of models with 3-OHK for the Y+ treatment. Mylar film resembles the UV filter but acts as a neutral-density filter. The red portions of the wings were identical in both Y+ and Y− treatments.

For the UV+ models, an odorless UV-reflective yellow paint (Fish Vision^TM^) was added to the dorsal and ventral yellow band of the model wings to provide UV reflectance (Fig. 1A,B, blue line), and the red portions of the wings were printed as described in Finkbeiner et al. (2014). For UV- models, a thin film UV filter was placed over both the yellow and red/pink UV- reflective portions on the wings. The UV filter prevents any light reflectance up to 400 nm (Fig. 1A-D, blue-green line). Mylar film was added to the yellow and red/pink portions of models used for the UV+ treatment to function as a control.

Reflectance spectra of the paper models and individual *Heliconius erato petiverana* (n=15), *Eueides isabella, E. surdus, E. thales* (n=3/species) and *E. heliconoides* (n=2) butterfly wings were measured by first aligning each measured wing in the same orientation as shown in appendix B of Bybee et al. (2012). If the viewer was looking directly from above at the oriented wings, the fixed probe holder (Ocean Optics RPH-1) was placed horizontally on top of the wing such that the axis of the illuminating and detecting bifurcating fiber (Ocean Optics R400-7-UV/VIS) was at an elevation of 45°to the plane of the wing and pointed left with respect to the body axis. Illumination was by a DH-2000 deuterium-halogen lamp, and reflectance spectra were measured with an Ocean Optics USB2000 spectrometer. A spectralon white standard (Ocean Optics WS-1) was used to calibrate the spectrometer. For the irradiance spectra measurements, the USB2000 spectrometer, a calibrated tungsten light source (Ocean Optics LS-1-CAL), a 100 or 400 μm diameter fiber (Ocean Optics P100‐ or P400-2-UV-Vis) and cosine corrector (Ocean Optics CC-3-UV), which produces vector irradiance measures, were used (Cronin et al., 2014). Five irradiance spectra measurements of down-dwelling light were taken and averaged per site.

For the mate choice experiments, the von Kries’ tranformed quantum catches for stimuli (Kelber et al., 2003) were first calculated for *H. erato* males and *H. erato* females separately using high light intensity and sunny cage irradiance spectra. Pairwise discriminabilities between artificial models and natural wing reflectance spectra were determined using a trichromatic vision model for *H. erato* males and tetrachromatic vision models for *H. erato* females, respectively (Vorobyev and Osorio, 1998). Parameters for the butterfly visual models were as follows: Weber fraction=0.05 (Koshitaka et al., 2008), photoreceptor peak sensitivities, *λ*_max_=355 nm (female only), 390 nm, 470 nm and 555 nm, and relative abundances of cones, VS=0.13, B=0.2, G=1 (male) or UV=0.09, VS=0.07, B=0.17, G=1(female)(McCulloch et al., 2016). For the predation experiments, von Kries’ transformed quantum catches for only ventral wing stimuli (since the butterflies were presented with their wings folded) were calculated using high light intensity and irradiance spectra from two of the four habitats where the models were placed: forest cover and forest edge. (The other two habitats, Pipeline Road and paved road, were found to have normalized spectra that were identical to forest cover). Discriminabilities between stimili were determined using tetrachromatic models of bird vision representing two types of avian visual system, the UV-(blue tit, *Parus caeruleus*) and violet-type (chicken, *Gallus gallus*) systems. For chicken, we used ocular media of Toomey et al. (2016) and behaviorally-determined parameters of Olsson et al. (2015), namely, a Weber fraction=0.06 for the L cone, and relative abundances of cones (VS=0.25, S=0.5, M=1, L=1). For the blue tit, we followed the work of Hart et al. (2000) including the effects of blue tit ocular media and used a Weber fraction=0.05 for the L cone, and relative abundances of cones (UV=0.37, S=0.7, M=0.99, L=1).

### Mate Preference Experiments

To test whether *Heliconius* 3-OHK yellow and UV serve as visual signals for conspecifics, mate preference experiments were carried out using insectary facilities in Gamboa, Panama from September 2013 through February 2014. Data were collected from 80 wild-caught *H. erato petiverana* butterflies: 40 males and 40 females. Each butterfly was introduced individually into experimental cages (2 m × 2 m × 2 m) and presented with one of two pairs of the artificial butterfly models: Y+ versus Y−, or UV+ versus UV-. The models were separated by 1 m and attached to an apparatus used to simulate flight (see Finkbeiner et al., 2014). Movies 2 and 3 in Supplementary Information show an example of female butterfly trials with Y (Movie S2) and UV (Movie S3) models. Individual butterflies experienced six five-minute trials – three five-minute trials with each of the two pairs. During trials two variables were recorded: 1) approaches, which consisted of flight unequivocally directed toward the model, and in which the butterfly came within 20 cm of the model, and 2) courtship events, which were classified as sustained hovering or circling behavior around the model (for examples see Videos 2 and 3 in Finkbeiner et al., 2014). Mate preference data were analyzed using a two-way ANOVA in R to examine the effects of model type and sex. Measurements of spectral irradiance (see above) were taken to provide quantitative information about the illumination conditions during the trials (Fig. S2).

### Predation Experiments

Previously we have shown (Finkbeiner et al., 2014) that avian predators differentially attack achromatic local form butterfly models compared to chromatic models as well as models that display non-local or color-switched patterns (Fig. S1). Here we tested whether avian predators would differentially attack local wing color form paper models where UV or yellow is manipulated. Predation experiments were completed in Panama at the Smithsonian Tropical Research Institute Gamboa field station and at selected forest sites in Soberanía National Park (including Pipeline Road), from June through September in 2013. Models were fitted with plasticine abdomens and tied to branches with thread to represent natural resting postures in the following habitat types: forest cover (15 sites), forest edge (17 sites), Pipeline Road (unpaved road with partial forest cover, 55 sites), and paved road with partial forest cover (13 sites). Examples of foliage cover in each of these habitat types, along with corresponding spectral irradiance measurements, are presented in Fig. S3. For the 3-OHK yellow pigment study, five artificial models of each treatment (Y+ and Y−) were randomly placed in 100 forest sites (Finkbeiner et al., 2014). The sites were separated ~250 meters to account for avian predator home range (home ranges described in Finkbeiner et al., 2012). There were 500 Y+ models and 500 Y− models for a total of 1000 models. The same methods were used for the UV study, using 500 UV+ models and 500 UV- models in non-overlapping sites from the Y+/− models.

The models remained at their sites for four days, and each model was examined for evidence of predation. A butterfly was considered attacked if damage to the abdomen and wings appeared in the form of beak marks and/or large indentations in the abdomen (for examples of attacked models see Finkbeiner et al., 2012; Finkbeiner et al., 2014). The attack response was modeled as a binomial variable (yes or no) dependent upon butterfly model type using a zero-inflated Poisson regression model, including sites as a random effect, in R with the ‘pscl’ package (Zeileis et al., 2008; R Development Core Team, 2010; Jackman, 2011). To examine whether forest light environment affected predator behavior, the same analysis was used to compare predation between model types in four main habitat types: forest cover, forest edge, Pipeline Road (unpaved road with partial forest cover), and paved road with partial forest cover.

### Fluorescence Experiments

To determine the possible contribution of 3-OHK fluorescence to its yellow coloration we measured the absorption, excitation, and emission spectra of 1.5 mg 3-hydroxy-DL-kynurenine (3-OHK) (Sigma-Aldrich, Catalog No. H1771) in 3 ml methanol (Fisher Chemicals, Optima LC/MS grade, Catalog No. A456-1). The resultant solution was diluted to an optical density OD=0.3 to get it within the linear range for fluorescence measurement (Dhami et al., 1995). The absorption spectrum of the pigment was measured with a Cary-50 spectrometer (Varian), while the emission and excitation spectra was acquired with a Cary Eclipse fluorimeter (Varian). Quantum yield was determined using Coumarin 500 (Exciton, Catalog No. 05000) as a reference. The reflectance spectrum measurements of *H. erato* wings were made using an Ocean Optics USB2000 spectrometer, a UV-cut off filter (Edmund Optics #39-426), a 150 W Xenon Arc lamp (which resembles daylight illumination), and spectralon white standard.

## Results

### Discriminabilities of Model Spectra and Real Wings

To test the hypothesis that our Y+ and UV+ paper models resembled real *H. erato* yellow wing colors, and that our Y− and UV- paper modeled resembled real *Eueides* yellow wing colors, we calculated pairwise discriminabilities between real wings and model spectra. We did so for the male and female *H. erato* visual system, and then for the UV- and violet-type avian visual systems. We found that for both male and female *H. erato* eyes, Y+ was an excellent match to *H. erato* dorsal and ventral yellows, and that Y− and UV- were excellent matches to *Eueides* dorsal and ventral yellows under high light illumination (Table 1, 66.7-100% of pairwise comparisons fell below 1 JND and 100% fell below 2 JNDs). This means that under lower light levels, model spectra would be an even better match to real wings. For the UV+ treatment, only ventral yellow was an excellent match to the *H. erato* ventral yellow for either *H. erato* sex. From this we conclude that the Y+ paper model bears a strong resemblance to real *H. erato* yellow wings and the Y− paper model bears a strong resemblance to real *Eueides* yellow wings for *H. erato* butterflies under the experimental illuminant conditions in which they were tested.

**Table 1.** Percentage of *H. erato* and *Eueides* wing colors compared to paper models with chromatic JND values <0.5, <1, <2 for male and female *H. erato* under high light, sunny cage illumination. n=15 *H. erato;* n=9 *Eueides* specimens measured.

|  |  | Percent below the threshold |  |  |  |  |  |  |  |
| --- | --- | --- | --- | --- | --- | --- | --- | --- | --- |
|  |  | Y+ |  | Y- |  | UV+ |  | UV- |  |
|  |  | Female | Male | Female | Male | Female | Male | Female | Male |
| Dorsal yellow | 0.5JND | 0.0 | 6.7 | 0.0 | 0.0 | 0.0 | 0.0 | 36.4 | 63.6 |
|  | 1JND | 86.7 | 100.0 | 81.8 | 9.1 | 0.0 | 0.0 | 100.0 | 100.0 |
|  | 2JND | 100.0 | 100.0 | 100.0 | 100.0 | 0.0 | 0.0 | 100.0 | 100.0 |
| Ventral yellow | 0.5JND | 13.3 | 0.0 | 0.0 | 55.6 | 33.3 | 33.3 | 0.0 | 11.1 |
|  | 1JND | 86.7 | 86.7 | 100.0 | 77.8 | 100.0 | 100.0 | 88.9 | 66.7 |
|  | 2JND | 100.0 | 100.0 | 100.0 | 100.0 | 100.0 | 100.0 | 100.0 | 100.0 |
| Dorsal red | 0.5JND | 0.0 | 0.0 | 0.0 | 0.0 | 0.0 | 0.0 | 0.0 | 0.0 |
|  | 1JND | 13.3 | 0.0 | 13.3 | 0.0 | 6.7 | 0.0 | 0.0 | 0.0 |
|  | 2JND | 86.7 | 46.7 | 86.7 | 46.7 | 86.7 | 46.7 | 93.3 | 80.0 |
| Ventral red | 0.5JND | 46.7 | 46.7 | 46.7 | 46.7 | 46.7 | 46.7 | 0.0 | 0.0 |
|  | 1JND | 46.7 | 46.7 | 46.7 | 46.7 | 46.7 | 46.7 | 0.0 | 0.0 |
|  | 2JND | 100.0 | 66.7 | 100.0 | 66.7 | 100.0 | 73.3 | 60.0 | 100.0 |

For the UV- and VS-type avian visual systems, the match between Y+ and UV+ and *H. erato* ventral yellow and between Y− and UV- *Eueides* ventral yellow was less good than if these same stimuli were viewed by the butterflies (Table 2). These results indicate that for birds at least, under forest shade or edge illumination, no pair of stimuli fully captured the spectral differences between *Heliconius* or *Eueides* yellow wing colors. All pairs of model spectra used in behavioral experiments, however, differed by >1JND for both birds and butterflies (except for Y+ vs. Y− for ventral yellow viewed through the male eye)(Table 3). This indicates that for both birds and butterflies, there was sufficient difference between the four model types to potentially elicit a behavioral response in the experiments described below.

**Table 2.**
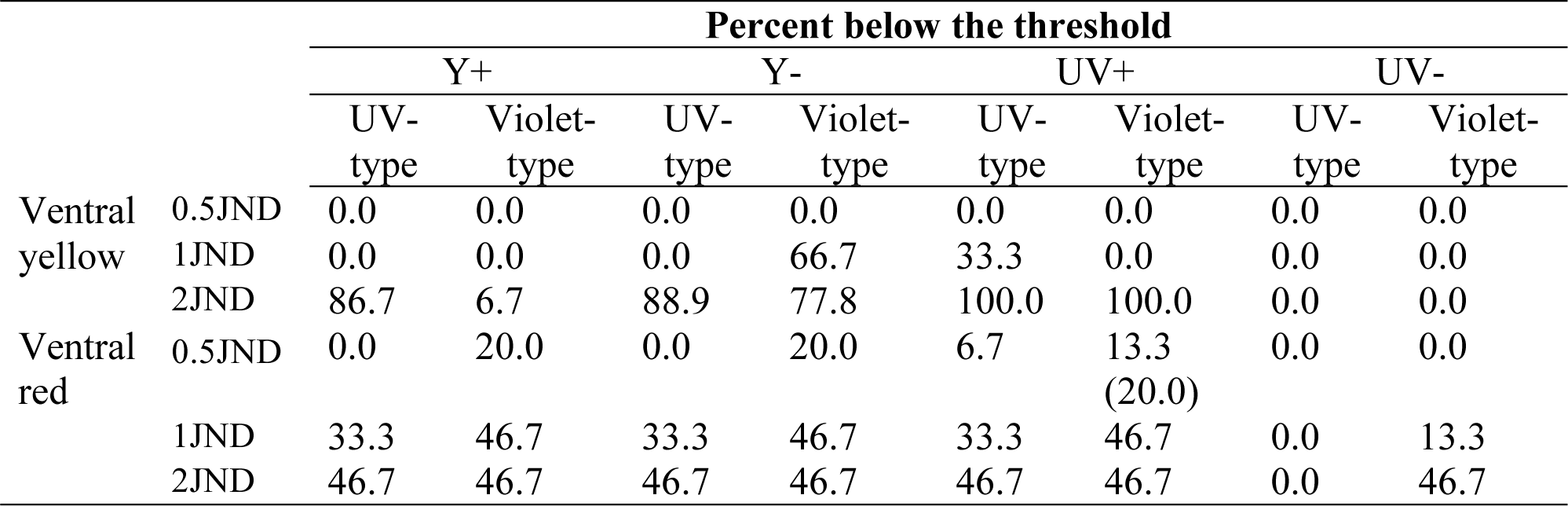
Percentage of *H. erato* and *Eueides* wing colors compared to paper models with chromatic JND values <0.5, <1, <2 for the UV-type, blue tit (*Parus caeruleus*) and violet-type chicken (*Gallus gallus*) under high light, partial forest shade illumination. n=15 *H. erato;* n=9 *Eueides* specimens measured. The percentages below the threshold were identical except for the number indicated in parentheses.

**Table 3.** JNDs between model spectra through the eyes of male and female *H. erato* and representatives of the UV‐ and violet-type bird visual systems. For butterflies, sunny cage illumination and for birds, partial forest cover illumination was used. Numbers in parentheses represent spectra modeled with forest edge illumination.

|  | JNDs |  |  |  |  |  |  |  |
| --- | --- | --- | --- | --- | --- | --- | --- | --- |
|  | Y+ vs. Y- |  |  |  | UV+ vs. UV+ |  |  |  |
|  | Butterfly |  | Bird |  | Butterfly |  | Bird |  |
|  | Female | Male | UV-type | VS-type | Female | Male | UV-type | VS-type |
| Dorsal yellow | 1.04 | 1.77 | N/A | N/A | 2.38 | 1.28 | N/A | N/A |
| Dorsal Red | N/A | N/A | N/A | N/A | 2.09 | 1.22 | N/A | N/A |
| Ventral yellow | 1.27 | 0.14 | 3.11<br>(3.37) | 1.86<br>(1.91) | 2.42 | 1.23 | 5.04<br>(5.38) | 0.97<br>(1.03) |
| Ventral Red | N/A | N/A | N/A | N/A | 2.28 | 1.23 | 4.73<br>(5.05) | 1.01 |

### Experiment 1: Effect of model type on mate preference

To determine how *Heliconius* yellow and UV affect conspecific recognition, we presented wild-caught *H. erato* butterflies with artificial butterfly models that had manipulated yellow and UV coloration. Preference toward models was measured in the form of approaches and courtship events. We found a strong model type effect on the number of butterfly approaches toward 3-OHK yellow and UV models. There were significantly more approaches toward Y+ than Y- models (Two-way ANOVA, F=16.287, p<0.0001, n=80), and toward UV+ than UV- models (F=10.469, p=0.002, n=80; Fig. 2A, black lines). There was no apparent effect of sex on butterfly approach behavior (F=2.738, p=0.099, n=80 for Y; F=0.049, p=0.952, n=80 for UV), suggesting that males and females approach the models at equal rates. Specific male and female behaviors for all comparisons are illustrated in Fig. S4.

**Fig. 2.**
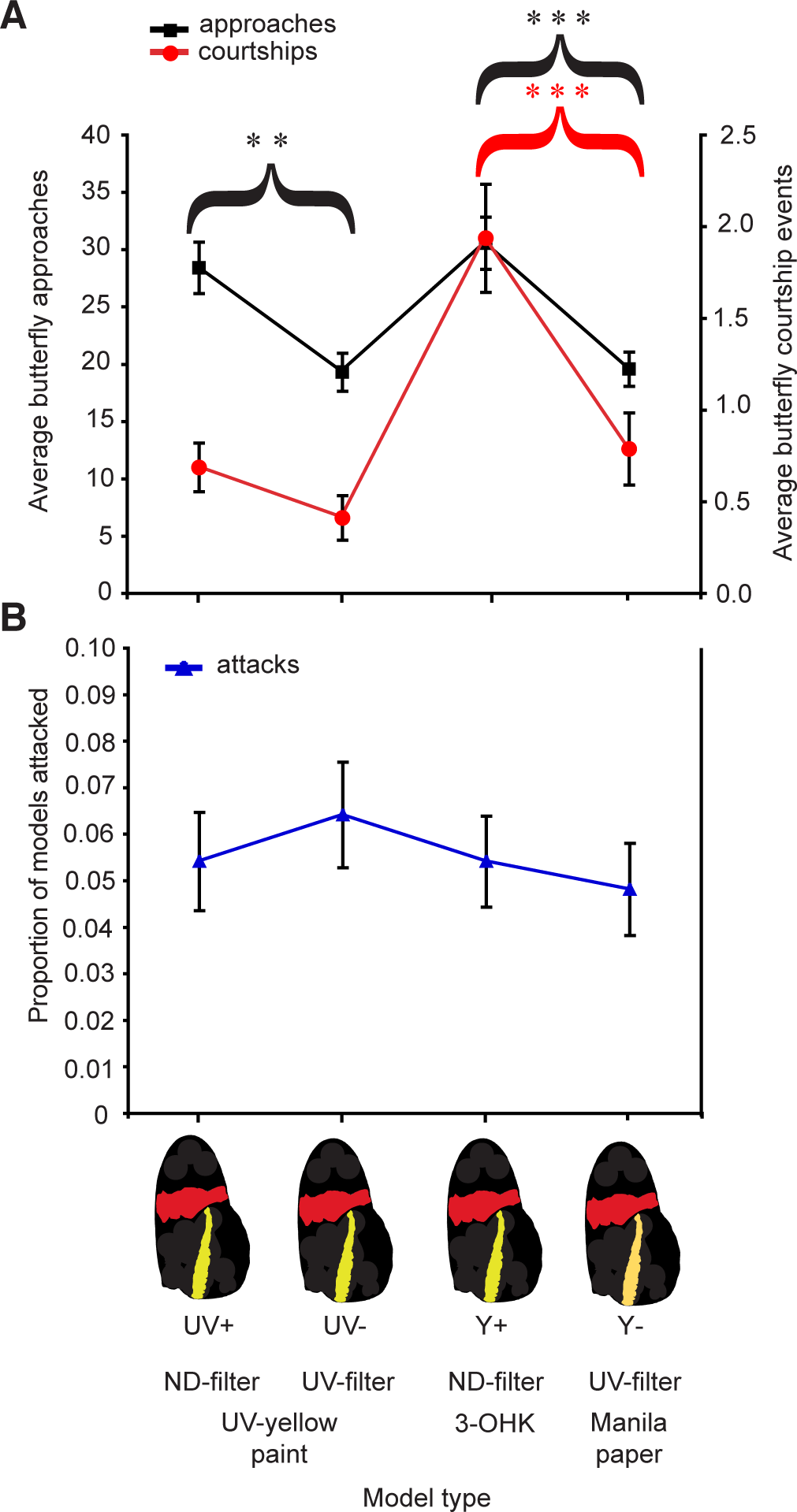
UV‐ and 3-OHK-manipulated butterfly models experience different rates of approach and courtship behavior by butterflies and similar rates of predation by birds. There are four model types that differ according to whether UV-yellow paint (UV+, UV-), 3-OHK pigment (Y+) or Manila paper (Y-) was used to produce the yellow hindwing bar and according to whether a neutral density filter (+ treatments) or a UV-blocking filter (− treatments) was used. (A) Mean approach (left axis black) and courtship (right axis, red) values with ±s.e.m. bars (each n = 80 butterflies: 40 males and 40 females). Asterisks represent the p-values from pairwise comparisons where *P<0.05, **P<0.01, ***P<0.001. (B) Average proportion of models attacked at each site (total n = 2000: 500 of each model type, 100 sites) with ±s.e.m. bars. The p-values from pairwise comparisons are >0.05.

Regarding courtship behavior, we found a strong model type effect where Y+ models were courted much more than Y- models (F=11.731, p=0.0008, n=80; Fig. 2A, red lines). The test for the main effect of sex shows that males court Y models at a significantly higher rate than females (F=9.211, p=0.0002, n=80). However, we found no significant model type effect on the number of courtship events directed toward UV+ and UV- models (F=2.304, p=0.131, n=80). There was also no effect of sex on butterfly courtship behavior toward the UV models (F=0.701, p=0.498, n=80).

### Experiment 2: Predator response to 3-OHK yellow and UV in different forest habitats

Previously we showed that birds preferentially attack achromatic *H. erato* models over Y+ chromatic models (Fig. S1) (Finkbeiner et al., 2014), as expected if chromatic cues serve as aposematic signals to avian predators. To test whether birds differentially attack UV- or yellow-manipulated models, predation was measured as the frequency of avian attacks on models in the forest. A total of 110 avian attacks were recorded (over four days of predator exposure for 500 models of each type): 27 and 24 attacks on Y+ and Y- models, and 27 and 32 attacks on UV+ and UV- models, respectively. Using a zero-inflated Poisson regression model, we detected no difference in predation between Y+ and Y- models: (z-value=-0.014, p=0.989, n=1000; Fig. 2B, blue lines), and no difference in predation between UV+ and UV- models: (z-value=-0.536, p=0.592, n=1000; Fig. 2B). A test of whether forest type affected predator behavior found no difference in predation between the model types in forest cover, forest edge, Pipeline Road (unpaved road with partial forest cover), and paved road with partial forest cover (all p-values >0.10). Although our prior experiments indicate that avian predators differentially attack *Heliconius erato* paper models that differ in both red and yellow color and pattern (Finkbeiner et al., 2014), the results presented here indicate that avian predators do not differentially attack 3-OHK yellow and other yellow or UV+ and UV- models in field trials.

### Fluorescence does not contribute to the yellow signal

The absorption spectrum of 3-OHK has a distinctive peak (*λ*_max_) at 380 nm (Fig. 3B), so this wavelength was chosen as the excitation wavelength for fluorescence measurements (10 nm bandwidth). The excitation spectrum of the pigment (Fig. 3C, black line) is in full agreement with absorption measurements demonstrating that the 380 nm is the peak excitation wavelength. The fluorescence of the pigment has a broad spectrum with peak of the emission around 508 nm (Fig. 3C, green line). Notably, the emission spectra of 3-OHK overlaps well with the visible portion of *Heliconius* yellow, suggesting the fluorescence of 3-OHK might in principle contribute to the signal in the visible range.

**Fig. 3.**
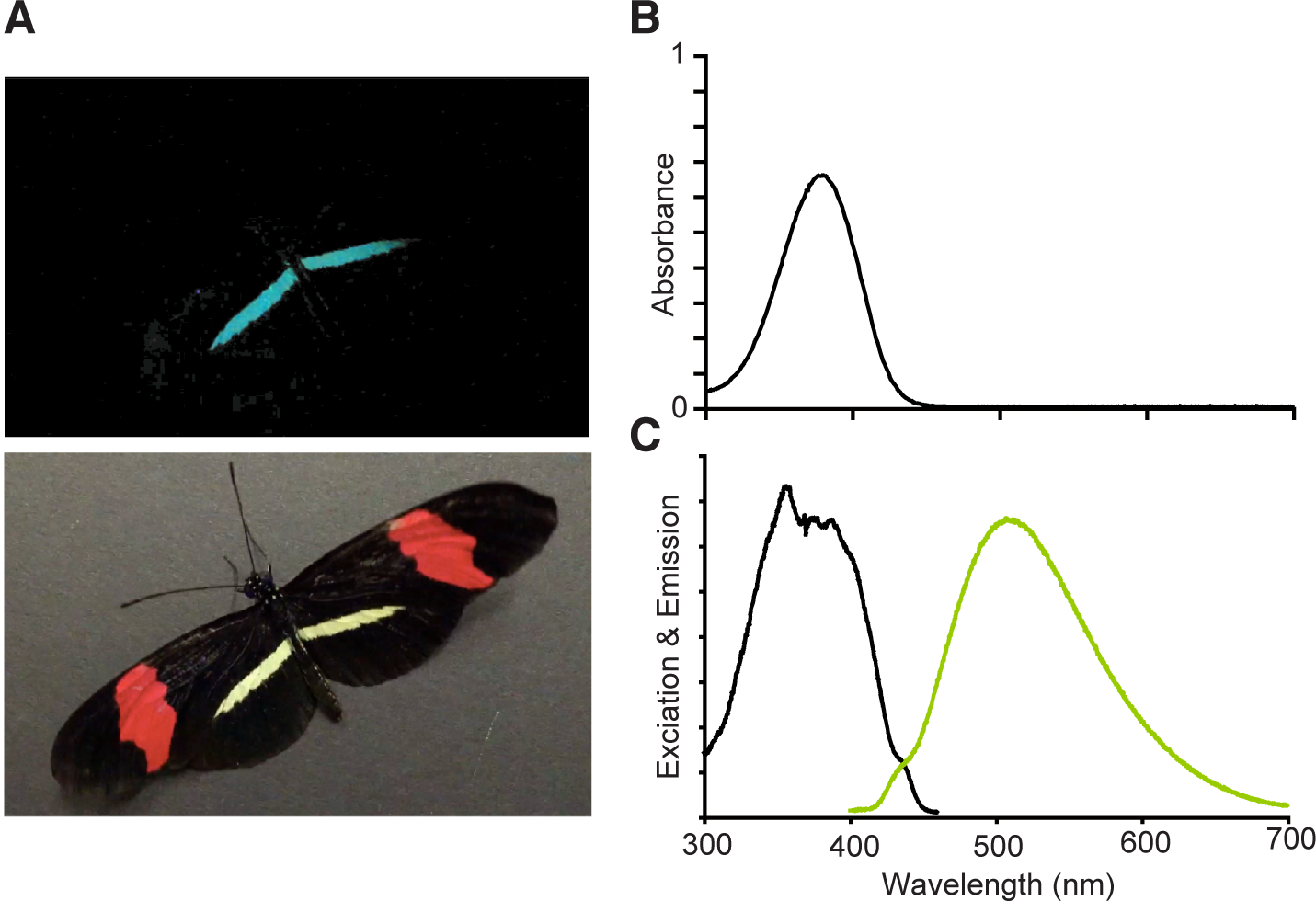
*Heliconius erato* fluorescence and 3-OHK absorption, excitation, and emission spectra. (A) Adult *H. erato* photographed under UV illumination to induce fluorescence (first panel) and under white light (second panel). (B) Absoprtion spectrum of 3-OHK in methanol (*λ*_max_=380 nm). (C) Excitation and emission spectrum of pigment 3-OHK. Emission has a broad spectrum with a peak around 508 nm.

In order to measure the efficiency of this emission, and hence understand if the fluorescence might contribute significantly to the signal, we determined the fluorescence quantum yield of 3-OHK. Quantum yield is characterized as the ratio of the number of photons emitted to the number of photons absorbed (Williams et al., 1983; Nad and Pal, 2003). Quantum yield was obtained by comparing 3-OHK to that of a standard and well-characterized fluorescent molecule, Coumarin 500 (Dhami et al., 1995), which has similar absorbance and fluorescence peaks as 3-OHK (Fig. S5). We were therefore surprised that the quantum yield of 3-OHK in methanol indicated that the emission is unlikely to be visible under normal illumination (quantum yield=5.1 x 10^−4^).

To be certain that these conclusions for 3-OHK in solution would also apply to 3-OHK on real wings in daylight illumination, additional experiments were carried out. Reflectance spectra of *H. erato* wings with and without a neutral-density filter (Mylar film) or a 400 nm cut-off filter (UV film), using a 150 W xenon arc lamp as a light source (which has a spectrum that resembles daylight illumination), were measured. If 3-OHK fluorescence does not contribute to the *Heliconius* yellow signal in broad-spectrum light, then measurements of *H. erato* wing reflectance spectra using a UV-cut off filter, which blocks excitation, should have no effect on the measured spectra in the visible range. That is indeed what we observed (Fig. S6). This series of experiments leads us to conclude that fluorescence does not contribute to the 3-OHK visual signal under broad-spectrum illumination.

## Discussion

### 3-OHK coloration is preferred by Heliconius erato

Butterflies are astonishingly diverse in their coloration, but the phylogenetic origins of new pigmentary coloration and the evolutionary forces that may have governed the adoption of a new pigment have rarely been investigated. Previously we showed that 3-OHK pigmentation is a synapomorphy of the genus *Heliconius*, being an ancestral character for the genus, but absent for sister genera such as *Eueides* (Briscoe et al., 2010). Here we have attempted to investigate how 3-OHK pigmentation functions as a signal for *H. erato* mate choice and defense. *Heliconius* yellow coloration has a spectrum, which includes reflectance maxima in the ultraviolet and human-visible range as well as fluorescence (Figs. 1A,B; 3A). Evidence here indicates that both the UV and long wavelength components of the reflectance spectrum contribute to the visual signal *H. erato* butterflies use for conspecific recognition, but qualitatively that the UV part may be less important for *H. erato* courtship than it is for approach behavior. Specifically the butterflies demonstrated clear preferences under all circumstances for Y+ over Y- (Fig. 2A). It is notable that our discriminability modeling of male and female *H. erato* vision indicates that for the butterflies at least the Y+ yellows are a good match to real *H. erato* yellow wing colors and Y- yellows are a good match to real *Eueides* yellow wing colors (Table 1). These results provide the first empirical evidence that *H. erato* butterflies prefer 3-OHK yellows to yellows found on the wings of their sister-genera, *Eueides*, and the first empirical evidence that the evolution of 3-OHK pigmentation in *Heliconius* may have been driven by sexual selection.

The interpretation of the UV+ and UV- treatments is a little less clear. Both UV+ and UV- models had the same long wavelength reflectance, but differed in the UV. UV+ models were approached by both sexes more frequently than UV- models, but while there was a trend towards preferring UV+ models during mating attempts, this difference was non-significant. This observation is perhaps surprising in view of the idea that at least for birds UV may be a short-range signal (Stevens and Cuthill, 2007). On the other hand, our discriminability calculations indicate that the UV+ dorsal yellow model color was not a good match to real *H. erato* dorsal yellow (Table 1). Neither the long wavelength nor the UV reflectance for dorsal yellow UV treatments were as similar to natural *H. erato* dorsal yellow as was the Y+ treatment (Fig. 1A, Table 1). It may be that a closer match to the natural *H. erato* spectrum—including in the UV—is needed to elicit a stronger courtship response.

Many prior studies of butterfly mate choice have examined the preferences of one sex or the other but not both (Knüttel and Fiedler, 2001; Fordyce et al., 2002; Ellers and Boggs, 2003; Sweeney et al., 2003; Kemp, 2007b). We note that our mate preference results indicate equal responses to models by males and females with respect to approach behavior. This shows that females are ‘active’ during such preference studies (see Movies S2 and S3), and that females and males may share similar preferences for *Heliconius* yellow and UV in conspecifics. In nature, females may use approach behavior in non-mating related interactions (Crane, 1955; Crane, 1957), such as following between pollen resources or to new roosting locations (Waller and Gilbert, 1982; Finkbeiner, 2014).

Our field study results show that 3-OHK yellow and UV do not alter avian predation rates in themselves, despite studies showing that birds use UV for mate recognition and foraging (Bennett et al., 1996; Siitari et al., 1999; Lyytinen et al., 2004). Recent work has shown that birds have even lower-than-expected UV sensitivity when looking at stimuli against a UV-poor background (Chavez et al., 2014) and understory-dwelling birds may have lower UV opsin expression than canopy-dwelling birds (Bloch, 2015). Our results resemble those of Lyytinen et al. (2000), who also found no support for UV as an aposematic signal for bird predators. Moreover we provide experimental evidence that in natural conditions, the mimicry between *Heliconius* yellow/UV coloration and non-*Heliconius* yellow/non-UV coloration in butterflies is successful for deterring birds. Given that we found no indication that *Heliconius* yellow and UV enhance aposematic signaling toward avian predators, this reinforces the notion that the phylogenetic switch from using other yellow pigments to 3-OHK as a signal on *Heliconius* wings is significant exclusively in relation to intraspecific communication.

### Fluorescence does not function as a signal

Several studies have concluded that fluorescence is an important component of complex signals in aquatic animals because of the contrast between narrow-band down-welling blue light and long-wavelength fluorescence (Mazel et al., 2004; Gerlach et al., 2014), but the evidence that fluorescence contributes to signaling in terrestrial animals, where the illumination spectrum is broad-band, is much more limited and somewhat mixed. For instance, one laboratory study of fluorescence in parakeets (*Melopsittacus undulatus*) suggested that fluorescence contributed to sexual signaling (Arnold et al., 2002) while two other studies of the same species did not (Pearn et al., 2001; Pearn et al., 2003). In spiders, lab studies indicate that fluorescence plays a role in male mate choice while UV plays a role in female mate choice (Lim et al., 2007). A paper investigating UV and fluorescence in damselfly signaling (Guillermo-Ferreira et al., 2014) concluded that there might be a possible contribution of fluorescence to the signal, however, important controls necessary to confirm this were absent.

To our knowledge, we report here for the first time that the yellow wing coloration of *Heliconius* is fluorescent (Fig. 3A); although Rawson (1968) mentions anecdotally that *H. erato* and *H. charithonia* wings are fluorescent but without specification whether it is the yellow portion of the wings, and without identification of the fluorescent chemical. We find by measuring the absorption, excitation, and emission spectrum and quantum yield of 3-OHK, together with wing reflectance spectra using daylight-simulating illumination, however, no evidence that 3-OHK fluorescence enhances the reflectance spectrum of *Heliconius* yellow under broad-band illumination. Although the spectral sensitivity of the *H. erato* blue receptor (470 nm) is well-suited to detecting 3-OHK fluorescence (McCulloch et al., 2016) we found no evidence that under natural illumination, fluorescence contributes to the 3-OHK signal in the visible range. Our result highlights the importance of quantifying fluorescence using several methods, and specifically under broad-band daylight-simulating illumination, before concluding that it contributes to a signal under terrestrial environments (e.g. Andrews et al., 2007).

## Conclusion

In summary, we demonstrate that *Heliconius* butterflies prefer 3-OHK yellow pigments in the context of conspecific signaling, these pigments have likely been selected for their reflectance properties in the visible range, and that fluorescence does not contribute to the visual signal. These results advance our understanding of the selective forces driving the transition from using other yellow pigments to using 3-OHK pigmentation in the genus *Heliconius*. We provide strong evidence that 3-OHK pigmentation is being maintained because it allows *Heliconius* species to recognize conspecifics for interspecific communication and sexual selection, whilst retaining the potential benefits of Müllerian mimicry with genera such as *Eueides*.

## Acknowledgements

We thank Robert Reed, Owen McMillan, Kailen Mooney, and Nancy Burley for advice and aid in project design; Ella Yuen, Angela Oh, Nina Chiu, Santiago Meneses, and David Carter for field assistance; Adriana Tapia, Oscar Paneso, and Elizabeth Evans for insectaries assistance; Brett Seymoure, Kyle McCulloch and Johannes Spaethe for assistance with taking irradiance and reflectance spectra; Darrell Kemp, Gil Smith, Kyle McCulloch, Jennifer Briner, Ana Catalán and Aide Macias-Muñoz for manuscript feedback; Dave Krueger and UCI ImageWorks for printing models; the Smithsonian Tropical Research Institute (STRI) for use of field sites; and La Autoridad Nacional del Ambiente (ANAM, Panama) for research permit approval.

## Author Contributions

S.D.F. designed butterfly models, carried out, and analyzed field predation and mate preference experiments, and wrote the manuscript; D.A.F. contributed measurements and analysis of physical fluorescent properties; D.O. and A.D.B. conceived of the study and edited the manuscript; A.D.B. designed butterfly models, calculated discriminabilities, performed experiments, analyzed fluorescence data and wrote the manuscript. All authors gave final approval for publication.

## Data Availability

DRYAD (doi: XXXXX)

## Supplementary Material Legends

**Movie S1:** Example of fluorescing 3-OHK pigment on a *H. erato* butterfly under a hand-held 365 nm LED light.

**Movie S2:** A female *H. erato* butterfly directs approaches toward a Y+ model (right side).

**Movie S3:** A female *H. erato* butterfly directs approaches toward a UV+ model (left side).

**Fig. S1.**
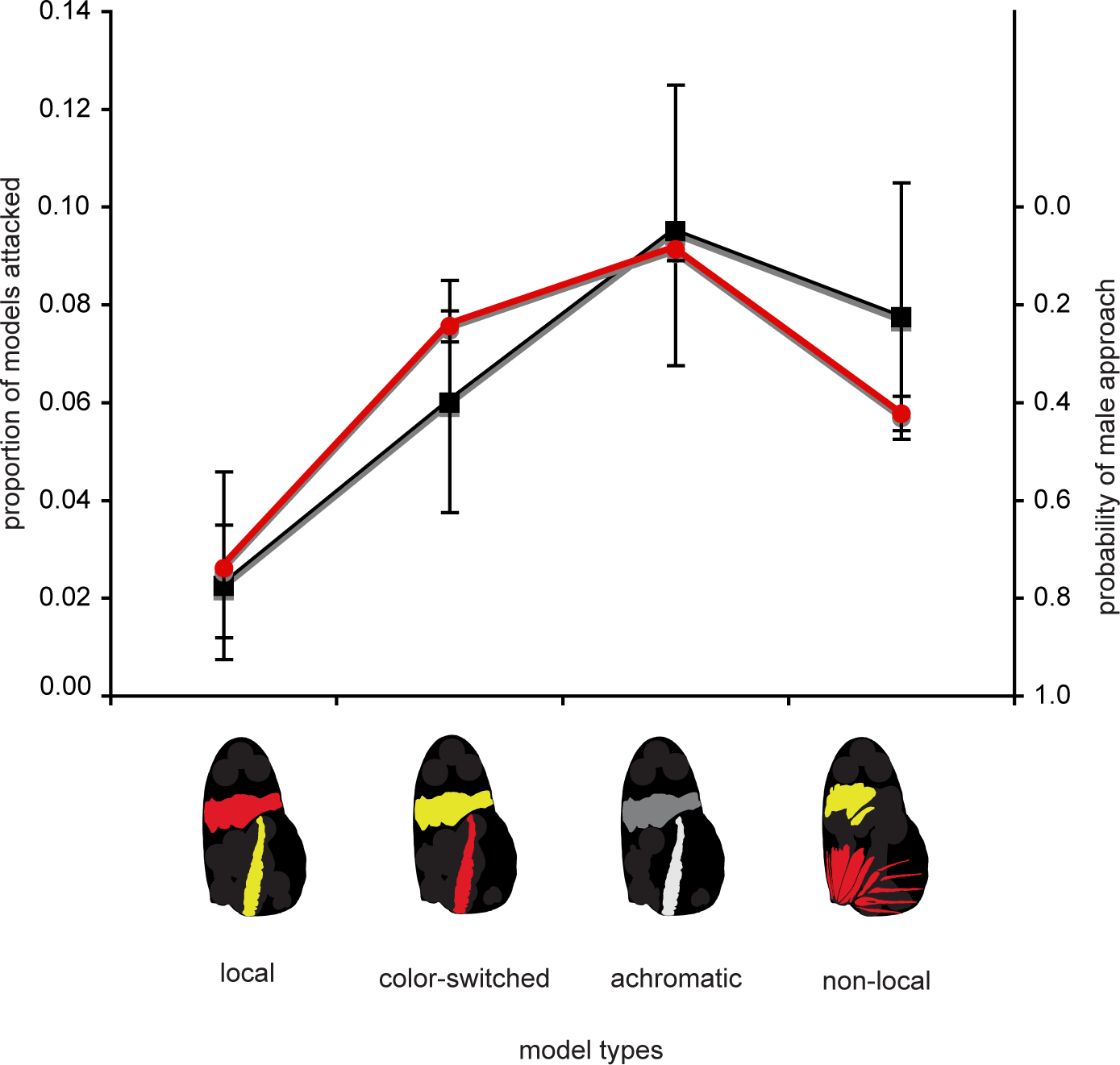
Color‐ and pattern-manipulated butterfly models experience different predation rates (left axis) and different probabilities of inducing premating approach behavior in male butterflies (right axis). There are four model types: a local *H. erato* type, a color-switched type, an achromatic type, and a nonlocal type. ±s.e.m. bars for the predation data include 95% CIs and ±s.e.m. bars for the mate preference data represent 95% credible intervals. Asterisks represent the p-values from pairwise comparisons between predation on the local model type and the three other model types where *P<0.05, **P<0.005, ***P<0.0001. All approach probability comparisons show that the preference means differsignificantly between the model types (Reprinted with permission from Finkbeiner et al. 2014).

**Fig. S2.**
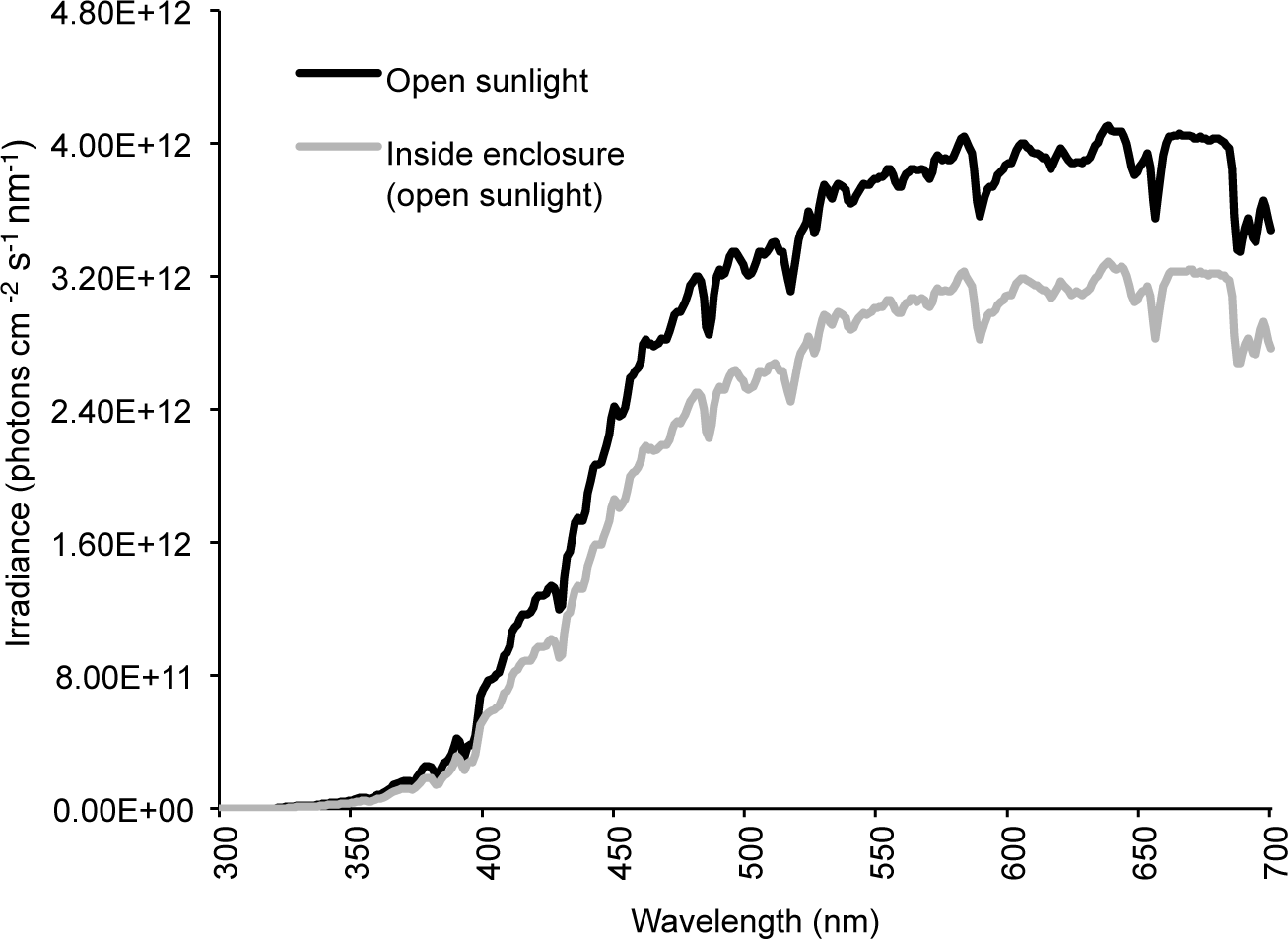
Irradiance spectra of open sunlight and the experimental cage during open sunlight conditions. Each graph represents the average from five measurements in each condition.

**Fig. S3.**
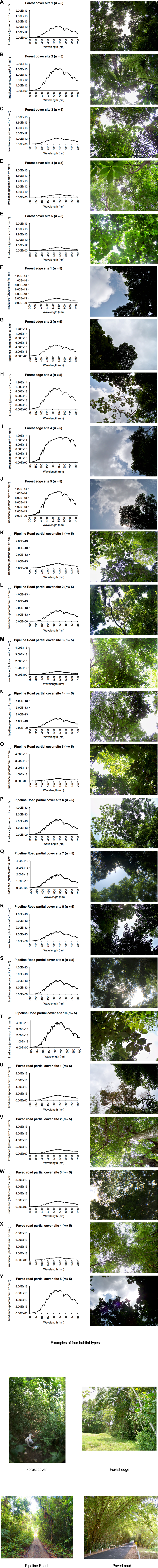
Habitat types. Irradiance spectra with photos of corresponding foliage cover, taken from the four major habitat types used in the predation study: forest cover (A-E); forest edge (J); Pipeline Road (unpaved road with partial forest cover), (K-T); and paved road with partial forest cover (U-Y). Five different sites were measured (repeated five times) for forest cover, forest edge, and paved road, whereas ten different sites were measured (repeated five times) for Pipeline Road because this was the dominant habitat type used in the study.

**Fig. S4.**
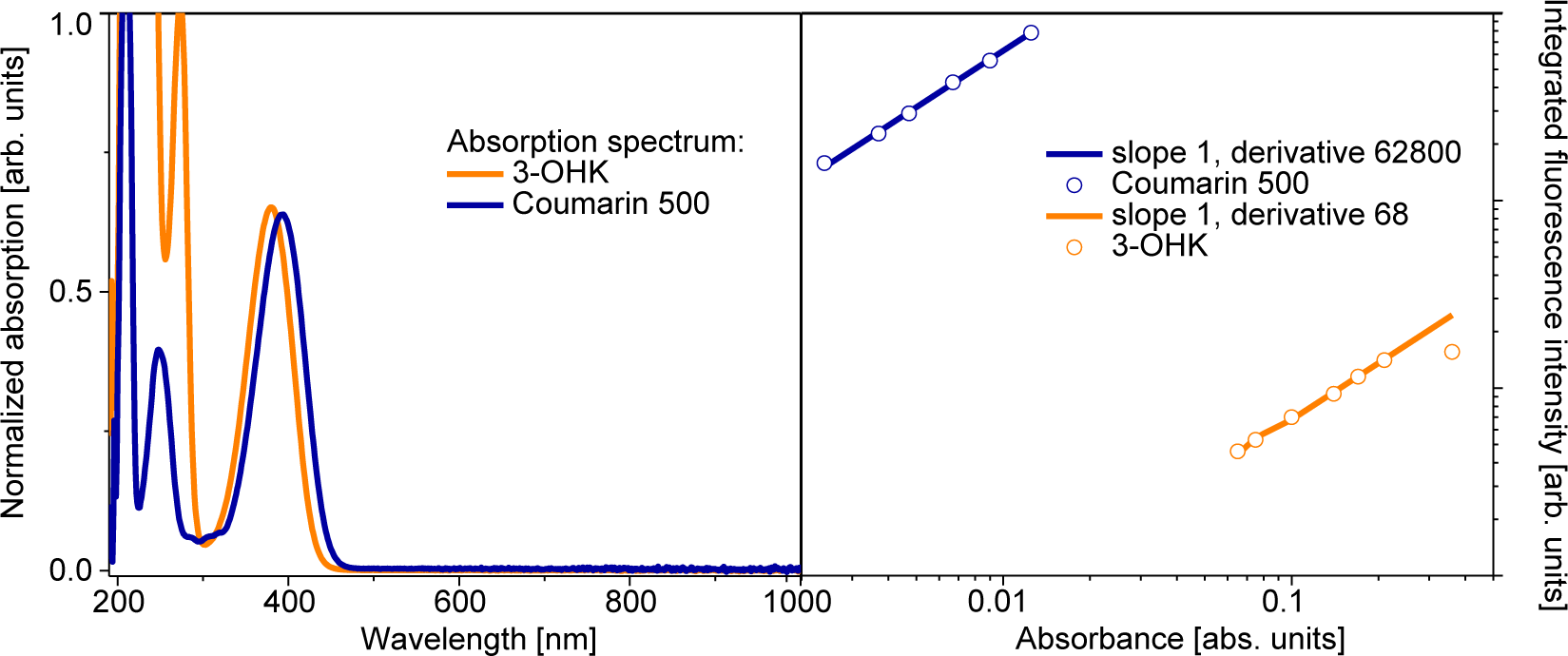
Male and female *H. erato* approach and courtship behavior. Male and female *H. erato* butterflies approach and court UV- and Y- manipulated artificial butterfly models at varying rates (A-D). All behaviors directed toward UV models are in the left column, and behaviors directed toward Y models are in the right column. Shown are the mean approach and courtship values ± s.e.m. (n=80 butterflies: 40 males and 40 females).

**Fig. S5.**
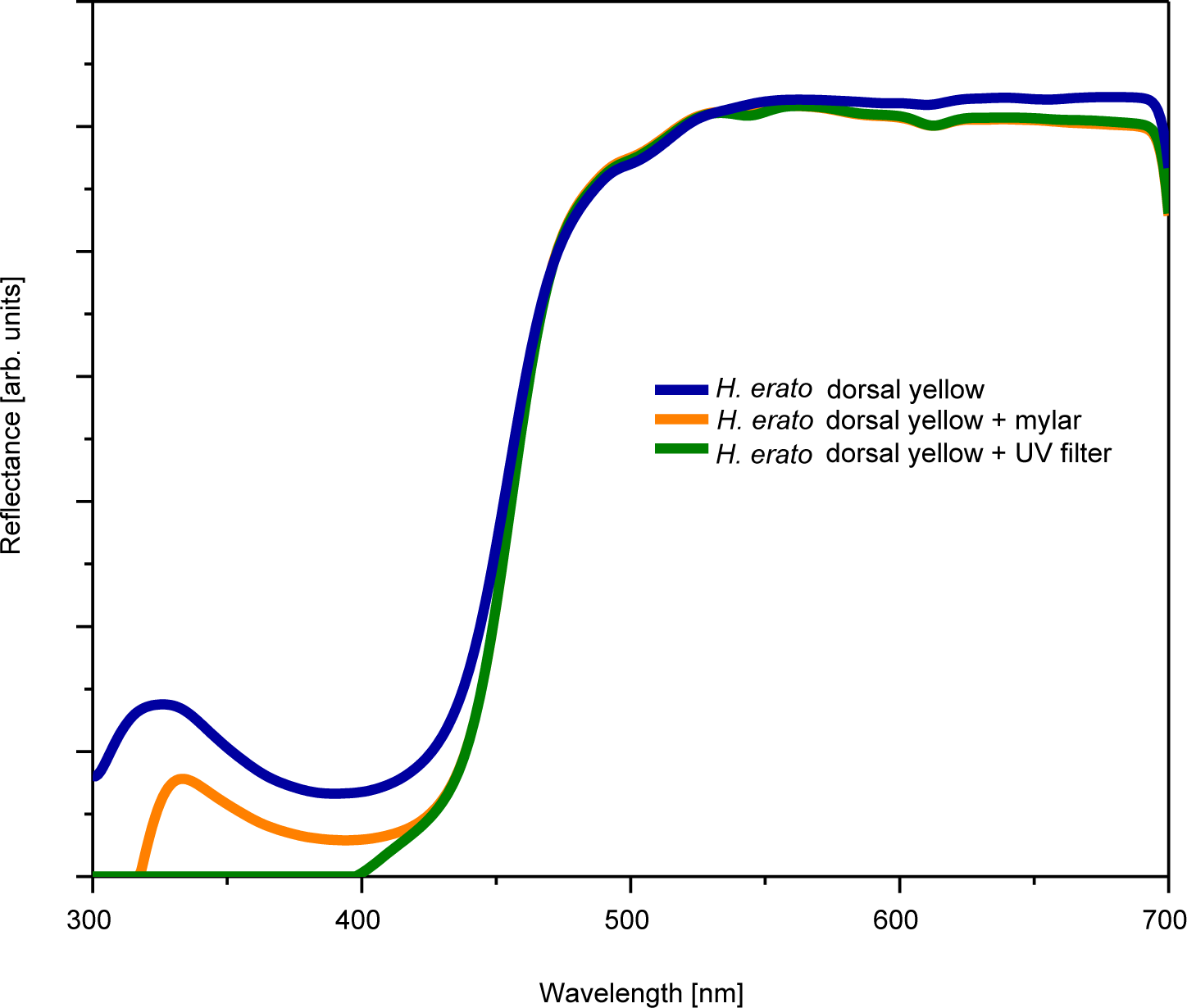
Experimental data used to determine the quantum yield of 3-OHK in methanol. (A) Absorption spectrum of 3-OHK pigment and Coumarin 500. Both dye and pigment have a very similar absorption spectrum making Coumarin 500 a good choice as a reference in quantum yield measurements. (B) Quantum yield determination using Coumarin 500 dye (blue curve) and 3-OHK pigment (orange curve). Coumarin 500 quantum yield is 0.46.

**Fig. S6.**
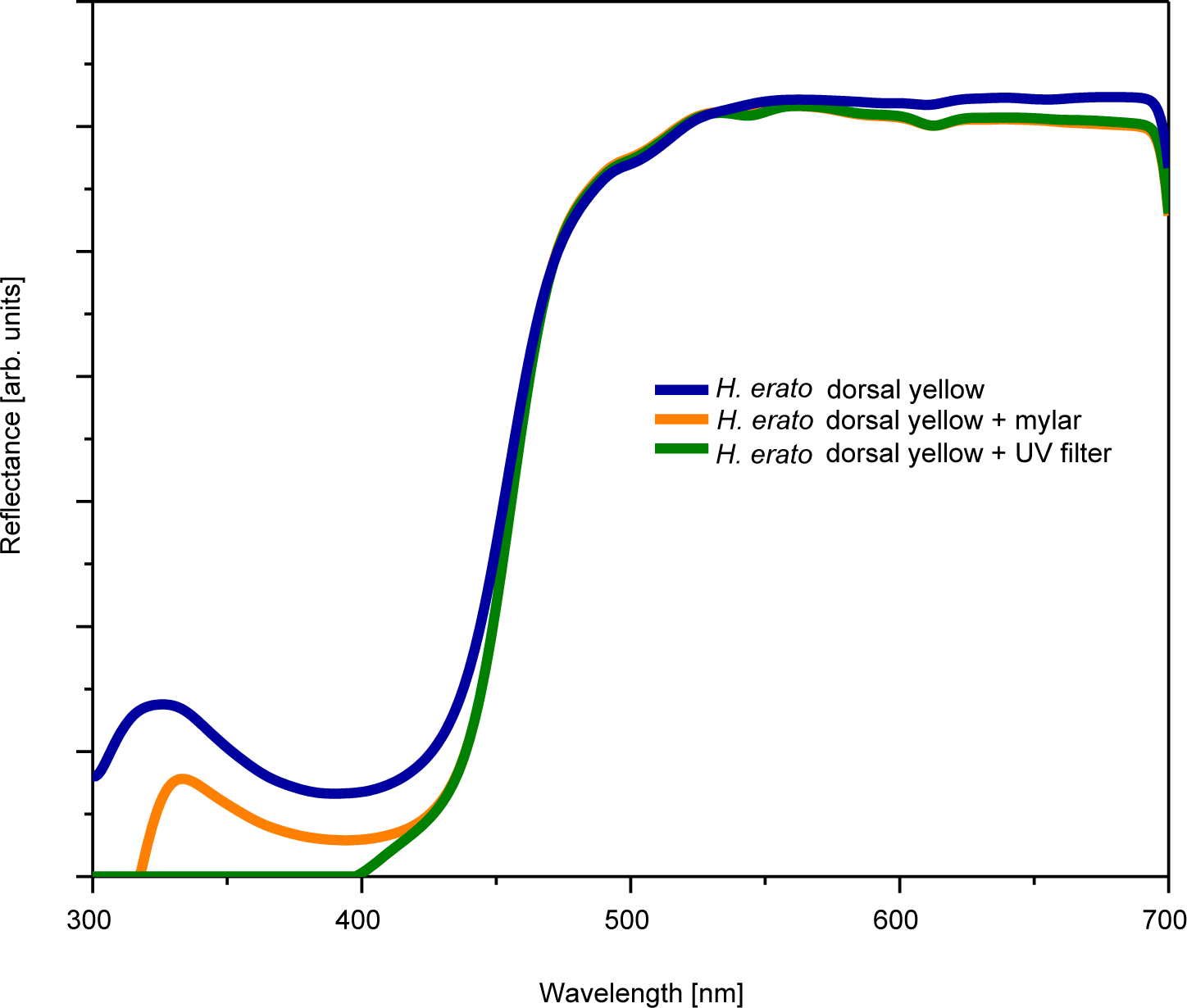
Reflectance spectrum of *H. erato* dorsal yellow hind wing with and without neutral density or UV-cutoff filters as measured using daylight-simulating illumination. The neutral density filter (Mylar) has an identical spectrum to the UV-cutoff filter in the visible range (above 400 nm) indicating that UV-induced fluorescence has no impact on the reflectance spectrum of 3-OHK yellow.

## References

Andrews, K., Reed, A. M. and Masta, S. E. (2007). Spiders fluoresce variably across many taxa. Biol. Lett. 3, 265–267. doi:10.1098/rsbl.2007.0016

Arikawa, K., Wakakuwa, M., Qiu, X., Kurasawa, M. and Stavenga, D. G. (2005). Sexual dimorphism of short-wavelength photoreceptors in the small white butterfly, Pieris rapae crucivora J. Neurosci. 25, 5935–5942. doi:10.1523/JNEUROSCI.1364-05.2005

Arnold, K. E., Owens, I. P. F. and Marshall, N. J. (2002). Fluorescent signaling in parrots. Science 295, 92. doi:10.1126/science.295.5552.92

Bennett, A. T. D., Cuthill, I. C., Partridge, J. C. and Maier, E. J. (1996). Ultraviolet vision and mate choice in zebra finches. Nature 380, 433–435. doi:10.1038/380433a0

Benson, W. W. (1972). Natural selection for Müllerian mimicry in Heliconius erato in Costa Rica. Science 176, 936–939. doi:10.1126/science.176.4037.936

Bloch, N. I. (2015). Evolution of opsin expression in birds driven by sexual selection and habitat. Proc. R. Soc. Lond. B. 282, 20142321. doi:10.1098/rspb.2014.2321

Briscoe, A. D. and Bernard, G. D. (2005). Eyeshine and spectral tuning of long wavelength-sensitive rhodopsins: No evidence for red-sensitive photoreceptors among five Nymphalini butterfly species. J. Exp. Biol. 208, 687–696. doi:10.1242/jeb.01453

Briscoe, A. D., Bybee, S. M., Bernard, G. D., Yuan, F., Sison-Mangus, M. P., Reed, R. D., Warren, A. D., Llorente-Bousquets, J. and Chiao, C.-C. (2010). Positive selection of a duplicated UV-sensitive visual pigment coincides with wing pigment evolution in Heliconius butterflies. Proc. Natl. Acad. Sci. USA. 107, 3628–3633. doi:10.1073/pnas.0910085107

Bybee, S. M., Yuan, F., Ramstetter, M. D., Llorente-Bousquets, J., Reed, R. D., Osorio, D. and Briscoe, A. D. (2012). UV photoreceptors and UV-yellow wing pigments in Heliconius butterflies allow a color signal to serve both mimicry and intraspecific communication. Am. Nat. 179, 38–51. doi:10.1086/663192

Chai, P. (1986). Field observations and feeding experiments on the responses of rufous-tailed jacamars (Galbula ruficauda) to freeflying butterflies in a tropical rainforest. Biol. J. Linn. Soc. 29, 161–189. doi:10.1111/j.1095-8312.1986.tb01772.x

Chavez, J., Kelber, A., Vorobyev, M. and Lind, O. 2014. Unexpectedly low UV-sensitivity in a bird, the budgerigar. Biol. Lett. 10, 20140670. doi:10.1098/rsbl.2014.0670

Chen, P.-J., Arikawa, K. and Yang, E.-C. (2013). Diversity of the photoreceptors and spectral opponnency in the compound eye of the golden birdwing, Troides aeacus formosanus. PLoS One 8,e62240. doi:10.1371/journal.pone.0062240.

Crane, J. 1955. Imaginal behavior of a Trinidad butterfly, Heliconius erato hydara Hewitson, with special reference to the social use of color. Zoologica 40, 167–196.

Crane J. 1957. Imaginal behavior in butterflies of the family Heliconiidae: Changing social patterns and irrelevant actions. Zoologica 42, 135–145.

Cronin, T. W. and Bok, M. J. (2016). Photoreception and vision in the ultraviolet. J. Exp. Biol. 219, 2790–2801. doi:10.1242/jeb.128769

Cronin, T. W., Johnsen, S., Marshall, N. J. and Warrant, E. J. (2014). Visual Ecology. Princeton, NJ, USA: Princeton University Press. 432 pp.

Cummings, M. E., Rosenthal, G. G. and Ryan, M. J. (2003). A private ultraviolet channel in visual communication. Proc. R. Soc. Lond. B. 270, 897–904. doi:10.1098/rspb.2003.2334

Dell’Aglio, D. D., Stevens, M. and Jiggins, C. D. (2016). Avoidance of an aposematically coloured butterfly by wild birds in a tropical forest. Ecol. Entomol. 41, 627–632. doi:10.1111/een.12335

Detto, T. and Blackwell, P.R.Y. (2009). The fiddler crab Uca mjoebergi uses ultraviolet cues in mate choice but not aggressive interactions. Anim. Behav. 78, 407–411. doi:10.1016/j.anbehav.2009.05.014

Dhami, S., de Mello, A. J., Rumbles, G., Bishop, S. M., Phillips, D. and Beeby, A. (1995). Phthalocyanine fluorescence at high concentration: dimers or reabsorption effect? Photochem. Photobiol. 61, 341. doi:10.1111/j.1751-1097.1995.tb08619.x

Eguchi, E. and Meyer-Rochow, V. B. (1983). Ultraviolet photography of forty-three species of Lepidoptera representing ten families. Annotationes Zoologicae Japoneses 56, 10–18.

Ellers, J. and Boggs, C. L. (2003). The evolution of wing color: male mate choice opposes adaptive wing color divergence in Colias butterflies. Evolution 57, 1100–1106. doi:10.1111/j.0014-3820.2003.tb00319.x

Finkbeiner, S. D. (2014). Communal roosting in Heliconius butterflies (Nymphalidae): Roost recruitment, establishment, fidelity, and resource use trends based on age and sex. J. Lep. Soc. 68, 10–16. doi:10.18473/lepi.v68i1.a2

Finkbeiner, S. D., Briscoe, A. D. and Reed, R. D. (2012). The benefit of being a social butterfly: communal roosting deters predation. Proc. R. Soc. Lond. B. 279, 2769–2776. doi:10.1098/rspb.2012.0203

Finkbeiner, S. D., Briscoe, A. D. and Reed, R. D. (2014). Warning signals are seductive: Relative contributions of color and pattern to predator avoidance and mate attraction in Heliconius butterflies. Evolution 68, 3410–3420. doi:10.1111/evo.12524

Fordyce, J. A., Nice, C. C., Forister, M. L. and Shapiro, A. M. (2002). The significance of wing pattern diversity in the Lycaenidae: mate discrimination by two recently diverged species. J. Evol. Biol. 15, 871–879. doi:10.1046/j.1420-9101.2002.00432.x

Gerlach, T., Sprenger, D. and Michiels, N. K. (2014). Fairy wrasses perceive and respond to their deep red fluorescent coloration. Proc. R. Soc. Lond. B. 281, pii: 20140787. doi:10.1098/rspb.2014.0787

Ghiradella, H. (1974). Development of ultraviolet-reflecting butterfly scales: how to make an interference filter. J. Morphol. 142, 395–410. doi:10.1002/jmor.1051420404

Grether, G. F., Kolluru, G. R. and Nersissian, K. (2004). Individual colour patches as multicomponent signals. Biol. Rev. 79, 583–610. doi:10.1017/S1464793103006390

Guillermo-Ferreira, R., Therézio, E. M., Gehlen, M. H., Bispo, P. C. and Marletta, A. (2014). The role of wing pigmentation, UV and fluorescence as signals in a neotropical damselfly. J. Insect Behav. 27, 67–80. doi:10.1007/s10905-013-9406-4

Hart, N. S., Partridge, J. C., Cuthill, I.C. and Bennett, A. T. D. (2000). Visual pigments, oil droplets, ocular media and cone photoreceptor distribution in two species of passerine bird: the blue tit (Parus caeruleus L.) and the blackbird (Turdus merula L.). J. Comp. Physiol. A. 186, 375–387. doi:10.1007/s003590050437

Jackman, S. (2011). pscl: Classes and methods for R developed in the political science computational laboratory, Stanford University. Department of Political Science, Stanford University. Stanford, California. R package version 1.04.1.

Johnsen, A., Andersson, S., Örnborg, J. and Lifjeld, J. T. (1998). Ultraviolet plumage ornamentation affects social mate choice and sperm competition in blue throats (Aves: Luscinia s. svecica): a field experiment. Proc. R. Soc. Lond. B. 265, 1313–1318. doi:10.1098/rspb.1998.0435

Kelber, A., Vorobyev, M. and Osorio, D. (2003). Animal colour vision: behavioural tests and physiological concepts. Biol. Rev. 78, 81–118. doi:10.1017/S1464793102005985

Kemp, D. J. (2007a). Female mating biases for bright ultraviolet iridescence in the butterfly Eurema hecubae (Pieridae). Behav. Ecol. 19, 1–8. doi:10.1093/beheco/arm094

------. Female butterflies prefer males bearing bright iridescent ornamentation. Proc. R. Soc. Lond. B. 274, 1043–1047. doi:10.1098/rspb.2006.0043

------. (2008). Female mating biases for bright ultraviolet iridescence in the butterfly Eurema hecabe (Pieridae). Behav. Ecol. 19, 1–8. doi:10.1093/beheco/arm094

Knüttel, H. and Fiedler, K. (2001). Host-plant-derived variation in ultraviolet wing patterns influences mate choice by male butterflies. J. Exp. Biol. 204, 2447–2459.

Koshitaka, H., Kinoshita, M., Vorobyev, M. and Arikawa, K. (2008). Tetrachromacy in a butterfly that has eight varieties of spectral receptors. Proc. R. Soc. Lond. B. 275, 947–954. doi:10.1371/journal.pone.0062240

Lagorio, M. G., Cordon, G. B. and Iriel, A. (2015). Reviewing the relevance of fluorescence in biological systems. Photochem. Photobio. Sci. 14, 1538–1559. doi:10.1039/C5PP00122F

Lim, L. M. L., Land, M. F. and Li, D. (2007). Sex-specific UV and fluorescence signals in jumping spiders. Science 315, 481. doi:10.1126/science.1134254

Llaurens, V., Joron, M., and Théry, M. (2014). Cryptic differences in colour among Müllerian mimics: how can the visual capacities of predators and prey shape the evolution of wing colours? J. Evol. Biol. 27, 531–540. doi:10.1111/jeb.12317

Lyytinen, A., Alatalo, R. V., Lindström, L. and Mappes, J. (2000). Can ultraviolet cues function as aposematic signals? Behav. Evol. 12, 65–70. doi:10.1093/oxfordjournals.beheco.a000380

Lyytinen, A., Lindström, L. and Mappes, J. (2004). Ultraviolet reflection and predation risk in diurnal and nocturnal Lepidoptera. Behav. Evol. 15, 982–987. doi:10.1093/beheco/arh102

Mazel, C. H., Cronin, T. W., Caldwell, R. L. and Marshall, N. J. (2004). Fluorescent enhancement of signaling in a mantis shrimp. Science 303, 51. doi:10.1126/science.1089803

McCulloch, K. J., Osorio, D. and Briscoe, A. D. (2016). Sexual dimorphism in the compound eye of Heliconius erato: a nymphalid butterfly with at least five spectral classes of photoreceptor. J. Exp. Biol. 219, 2377–2387. doi:10.1242/jeb.136523

Meyer-Rochow, V. B. (1991). Differences in ultraviolet wing patterns in the New Zealand lycaenid butterflies Lycaena salustius, L. rauparaha, and L. feredayi as a likely isolating mechanism. J. Roy. Soc. New Zealand 21, 169–177. doi:10.1080/03036758.1991.10431405

Nad, S. and Pal, H. (2003). Photophysical properties of Coumarin-500 (C500): Unusual behavior in nonpolar solvents. J. Phys. Chem. A. 107, 501–507. doi:10.1021/jp021141l

Obara, Y., Koshitaka, H. and Arikawa, K. (2008). Better mate in the shade: enhancement of male mating behaviour in the cabbage butterfly, Pieris rapae crucivora, in a UV-rich environment. J. Exp. Biol. 211, 3698–3702. doi:10.1242/jeb.021980

Olsson, P., Lind, O. and Kelber, A. (2015). Bird colour vision: behavioral thresholds reveal receptor noise. J. Exp. Biol. 218, 184–193. doi:10.1242/jeb.11187

Painting, C. J., Rajamohan, G., Chen, Z., Zeng, H., and Li, D. (2016). It takes two peaks to tango: the importance of UVB and UVA in sexual signalling in jumping spiders. Animal Behav. 113, 137–146. doi:10.1016/j.anbehav.2015.12.030

Pearn, S. M., Bennett, A. T. D. and Cuthill, I. C. (2001). Ultraviolet vision, fluorescence and mate choice in a parrot, the budgerigar Melopsittacus undulatus Proc. R. Soc. Lond. B. 268, 2273–2279. doi:10.1098/rspb.2001.1813

Pearn, S. M., Bennett, A. T. D. and Cuthill, I. C. (2003). The role of ultraviolet reflectance and ultraviolet-A induced fluorescence in the appearance of bugerigar plumage: insights from spectrofluorometry and reflectance spectrophotometry. Proc. R. Soc. Lond. B. 270, 859–865. doi:10.1098/rspb.2002.2315

Prum, R. O. and Torres, R. H. (2003). Structural colouration of avian skin: convergent evolution of coherently scattering dermal collagen arrays. J. Exp. Biol. 206, 2409–2429. doi:10.1242/jeb.00431

R Development Core Team. (2010). R: A language and environment for statistical computing. R Foundation for Statistical Computing, Vienna, Austria.

Rawson, G. W. (1968). Study of fluorescent pigments in Lepidoptera by means of paper partition chromatography. J. Lep. Soc. 22, 27–40.

Reed, R. D., McMillan, W. O. and Nagy, L. M. (2008). Gene expression underlying adaptive variation in Heliconius wing patterns: non-modular regulation of overlapping cinnabar and vermillion prepatterns. Proc. R. Soc. Lond. B. 275, 37–45. doi:10.1098/rspb.2007.1115

Robertson, K. A. and Monteiro, A. (2005). Female Bicyclus anynana butterflies choose males on the basis of their dorsal UV-reflective eyespot pupils. Proc. R. Soc. Lond. B. 272, 1541–1546. doi:10.1098/rspb.2005.3142

Rutowski R.L. (1977). The use of visual cues in sexual and species discrimination by males of the small sulphur butterfly Eurema lisa (Lepidoptera, Pieridae). J. Comp. Physiol. 115, 61–74. doi:10.1007/BF00667785

Rutowski, R. L., Macedonia, J. M., Morehouse, N. and Taylor-Taft, L. (2005). Pterin pigments amplify iridescent ultraviolet signal in males of the orange sulphur butterfly, Colias eurytheme. Proc. R. Soc. Lond. B. 272, 2329–2335. doi:10.1098/rspb.2005.3216

Shawkey, M. D. and Hill, G. E. (2005). Carotenoids need structural colours to shine. Biol. Lett. 1, 121–124. doi:10.1098/rsbl.2004.0289.

Siitari, H., Honkava, J. and Viitala, J. (1999). Ultraviolet reflection of berries attracts foraging birds. A laboratory study with redwings (Turdus iliacus) and bilberries (Vaccinium myrtillus). Proc. R. Soc. Lond. B. 266, 2125–2129. doi:10.1098/rspb.1999.0897

Silberglied, R. E. and Taylor, O. R. (1978). Ultraviolet reflection and its behavioral role in the courtship of the sulphur butterflies Colias eurytheme and C. philodice (Lepidoptera, Pieridae). Behav. Ecol. Sociobiol. 3, 203–243. doi:10.1007/BF00296311

Sison-Mangus, M. P., Briscoe, A. D., Zaccardi, G., Knüttel, H. and Kelber, A. (2008). The lycaenid butterfly Polyommatus icarus uses a duplicated blue opsin to see green. J. Exp. Biol. 211, 361–369. doi:10.1242/jeb.012617

Smith, E. J., Partridge, J. C., Parsons, K. N., White, E. M., Cuthill, I. C., Bennett, A. T. D. and Church, S. C. (2002). Ultraviolet vision and mate choice in the guppy (Poecilia reticulata). Behav. Ecol. 13, 11–19. doi:10.1093/beheco/13.1.11

Stalleicken, J., Labhart, T. and Mouritsen, H. (2006). Physiological characterization of the compound eye in monarch butterflies with focus on the dorsal rim area. J. Comp. Physiol. A. 192, 321–331. doi:10.1007/s00359-005-0073-6

Stavenga, D. G., Leertouwer, H. L., Marshall, N. J. and Osorio, D. (2011). Dramatic colour changes in a bird of paradise caused by uniquely structured breast feather barbules. Proc. R. Soc. Lond. B. 278, 2098–2104. doi:10.1098/rspb.2010.2293

Stavenga, D. G., Leertouwer, H. L. and Wilts, B. D. (2014). Coloration principles of nymphaline butterflies - thin films, melanin, ommochromes and wing scale stacking. J. Exp. Biol. 217, 2171–2180. doi:10.1242/jeb.098673

Stavenga, D. G., Stowe, S., Siebke, K., Zeil, J. and Arikawa, K. (2004). Butterfly wing colours: scale beads make white pierid wings brighter. Proc. R. Soc. Lond. B. 271, 1577–1584. doi:1098/rspb.2004.2781

Stevens, M. and Cuthill, I. C. (2007). Hidden Messages: Are ultraviolet signals a special channel in avian communication? BioScience 57, 501–507. doi:10.1641/B570607

Sweeney, A., Jiggins, C. and Johnsen, S. (2003). Polarized light as a butterfly mating signal. Nature 423, 31–32. doi:10.1038/423031a

Tokuyama, T. S., Senoh, S., Sakan, T., Brown, K. S. and Witkop, B. (1967). The photoreduction of kynurenic acid to kynurenine yellow and the occurrence of 3-hydroxy-L-kynurenine in butterflies. J. Am. Chem. Soc. 89, 1017–1021. doi:10.1021/ja00980a046

Toomey, M. B., Lind, O., Frederiksen, R., Curley, R. W., Riedl, K. M., Wilby, D., Schwartz, S. J., Witt, C. C., Harrison, E. A., Roberts, N. W. et al. (2016). Complementary shifts in photoreceptor spectral tuning unlock the full adaptive potential of ultraviolet vision in birds. eLife. 5,e15675.

Vorobyev, M. and Osorio, D. (1998). Receptor noise as a determinant of colour thresholds. Proc. R. Soc. Lond. B. 265, 351–358. doi:10.1098/rspb.1998.0302

Vukusic, P. and Hooper, I. (2005). Directionally controlled fluorescence emission in butterflies. Science 310, 1151. doi:10.1126/science.1116612

Vukusic, P. and Sambles, J. R. (2003). Photonic structures in biology. Nature 424, 853–855. doi: 10.1038/nature01941

Waller, D. A. and Gilbert, L. E. (1982). Roost recruitment and resource utilization: observations on Heliconius charithonia. J. Lep. Soc. 36, 178–184.

Watt, W. B. (1964). Pteridine components of wing pigmentation in the butterfly Colias eurytheme. Nature 201, 1326–1327. doi:10.1038/2011326b0

Williams, A. T. R., Winfield, S. A. and Miller, J. N. (1983). Relative fluorescence quantum yields using a computer controlled luminescence spectrometer. Analyst 108, 1067. doi:10.1039/AN9830801067

Yuan, F., Bernard, G. D., Le, J. and Briscoe, A. D. (2010). Contrasting modes of evolution of the visual pigments in Heliconius butterflies. Mol. Biol. & Evol. 27, 2392–2405. doi:10.1093/molbev/msq124

Zeileis, A., Kleiber, C. and Jackman, S. (2008). Regression Models for Count Data in R. J. Stat. Softw. 27, 1–25.

